# Exposure to PM_2.5_ during pregnancy or lactation increases methylation while reducing the expression of *Pdx1* and *NEUROG3* in mouse pancreatic islets

**DOI:** 10.1101/852509

**Authors:** Raquel Patricia Ataíde Lima, Vitor Ferreira Boico, Guilherme Francisco Peruca, Kellen Cristina da Cruz Rodrigues, Victor Yuji Yariwake, Daisuke Hayashi Neto, Maria José de Carvalho Costa, Junia Carolina Rebelo Santos-Silva, Gabriel Forato Anhê, Paulo Hilário Nascimento Saldiva, Mariana Matera Veras, Patrícia de Oliveira Prada

## Abstract

Air pollution is comprised of several substances, including particulate matter (PM). Exposure to air pollution may trigger alterations in DNA methylation thus modifying gene expression patterns. This phenomenon is likely to mediate the relationship between exposure to air pollution and adverse health effects. The purpose of this study was analyzing the effects of exposure to PM_2.5_ during pregnancy or lactation and whether it would cause multigenerational epigenetic alterations in the promoter region of the genes *Pdx1* and *NEUROG3* within mouse pancreatic islets. Our results show that maternal exposure to PM_2.5_ led to an elevation in blood glucose levels within the two following generations (F1 and F2). There was also an increase in DNA methylation in the aforementioned promoter regions accompanied by reduced gene expression in generations F1 and F2 upon F0 exposure to PM_2.5_ during pregnancy. These data suggest that maternal exposure to PM_2.5_ from air pollution, particularly during pregnancy, may lead to a multigenerational and lifelong negative impact on glucose homeostasis mediated by an increase in DNA methylation within the promoter region of the genes *Pdx1* and *NEUROG3* in pancreatic islets.

## 1. INTRODUCTION

Air pollution is a matter of utmost concern across the globe, intensified by modern phenomena such as rapid population growth, urban sprawl and nonstop increase in industrialization, electricity demand, deforestation, and the widespread use of motor vehicles moved by the combustion of fossil fuel [1].

Air pollution is comprised of several substances, mainly particulate matter (PM), which includes both liquid and solid air-suspended particles and is classified according to its source and size. Exposure to elevated levels of PM with a diameter ≤2.5 μm (PM_2.5_) has been associated with an increased risk for atherosclerosis, hypertension, cardiovascular diseases, and diabetes [2–4].

PM concentrations are particularly elevated in some developing countries, where they were shown to be well above the upper limit recommended by the World Health Organization’s guidelines [5], thus being regarded as a hazard to the health and the quality of life of people living in such areas [1,6].

Exposure to air pollution may lead to molecular alterations in genes, such as epigenetic modifications. Those alterations are capable of influencing health outcomes during all of the stages of life as well as of being inherited [7], thus potentially resulting in the transmission of new phenotypes across several generations [8].

DNA methylation is currently the most widely investigated epigenetic mechanism and occurs through the addition of a methyl group to a cytosine-guanine dinucleotide (CpG site) [9,10]. Alteration in DNA methylation may be induced by exposure to air pollution, thus altering gene expression patterns [11]. Those changes in DNA methylation are a plausible mechanism that potentially mediates the harm that air pollution causes to health.

Epidemiological and animal studies suggest that an adverse environment during early life, both intrauterine and postnatal, possibly increases the risk for type 2 diabetes [12]. Likewise, the long-term effects of exposure to these environments may be carried over to the following generations, affecting the offspring’s phenotype [13].

In this study, we exposed two groups of F0 generation mice (pregnant or lactating) to either filtered air or polluted air (PM_2.5_). F1 generation mice were exposed during either intrauterine development or within the first days after birth whereas F2 generation animals were potentially exposed through germ cells produced by F1 generation animals. The effects observed in the F1 and F2 generations are called “multigenerational”[14].

We examined the effects of maternal exposure to polluted air (PM_2.5_) during either pregnancy or lactation on F1 and F2 generations, and explored whether they were associated with epigenetic alterations in promoter regions of the genes *Pdx1* and *NEUROG3* in mouse pancreatic islets as an inherited epigenetic alteration.

## 2. RESULTS

### 2.1. Effects of PM_2.5_ exposure during either pregnancy or lactation on the following generation’s glucose homeostasis

In order to assess the impacts of exposure to polluted air on blood glucose levels and whether those effects could be inherited, 11-week old offspring of mice exposed to PM_2.5_ during either pregnancy or lactation had their fasting blood glucose measured. Both groups showed elevated results when compared to the control group, though only the offspring of animals exposed during lactation showed a significant difference when compared to those exposed to FA, with an average fasting blood glucose level of 101.6 mg/dL (PM_2.5_) and 90.2 mg/dL (FA) (*p* =0.0208) (figure 1, A and B). Fasting blood glucose remained elevated in the following generation (F2), with a significant difference among the group exposed to PM_2.5_ during lactation with an average fasting blood glucose level of 103.8 mg/dL (PM_2.5_) and 97 mg/dL (FA) (p = 0.0202) (figure 2, A and B).

**Figure 1.**
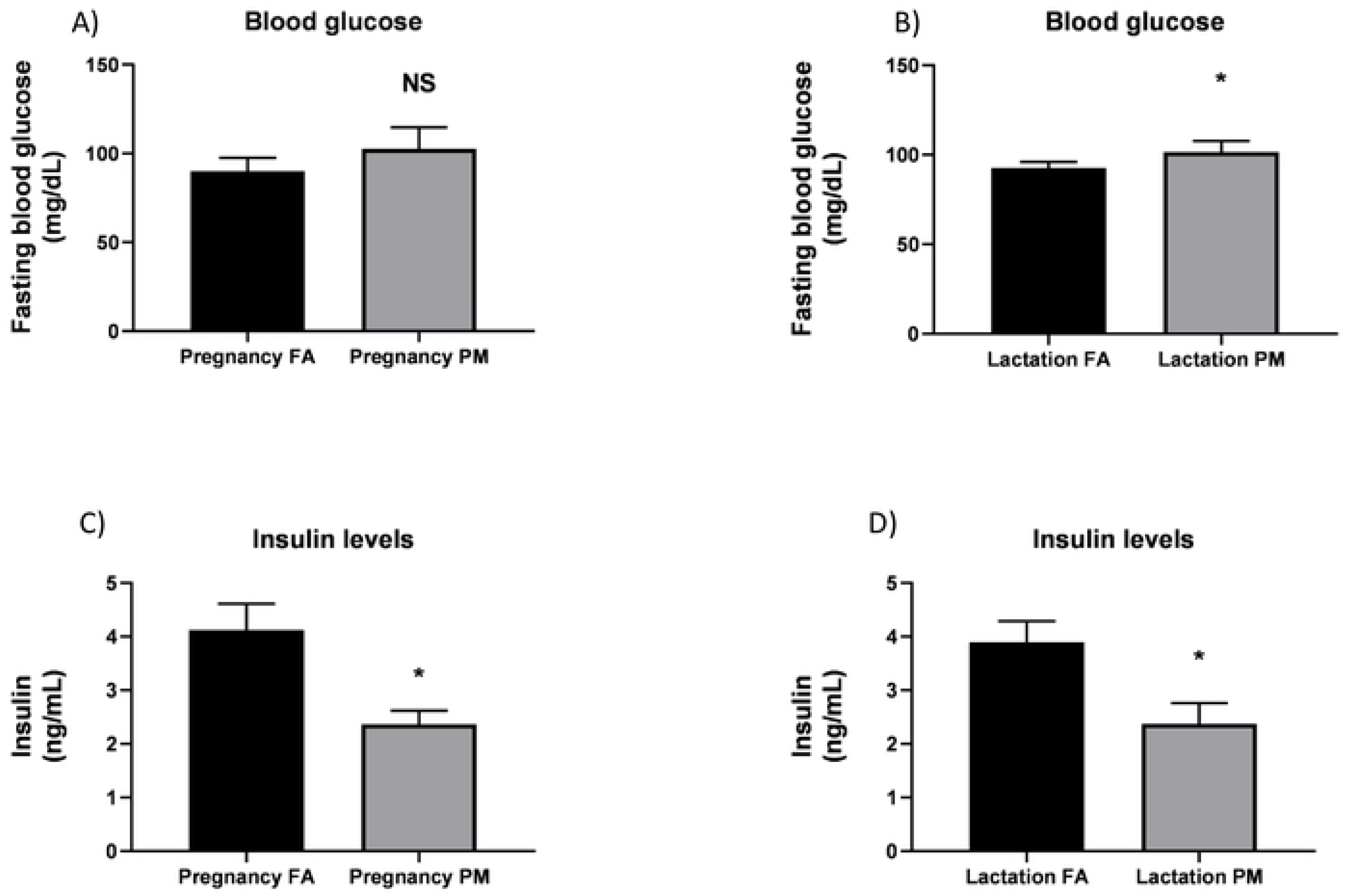
Blood glucose and insulin levels of the offspring (F1) of mice exposed to PM_2.5_ or FA during either pregnancy or lactation. Dams were exposed to PM_2.5_ (600 µg / m^3^ / day) or filtered air. Values include fasting blood glucose (mg/dL) and serum insulin (ng/mL). Data are shown as mean ± SD (n = 6 mice/group). Mice were 11-week old and fasted overnight before the experiments. A two-tailed Student’s t-test was used for statistical analysis. * *p* < 0.05 vs. FA.

**Figure 2.**
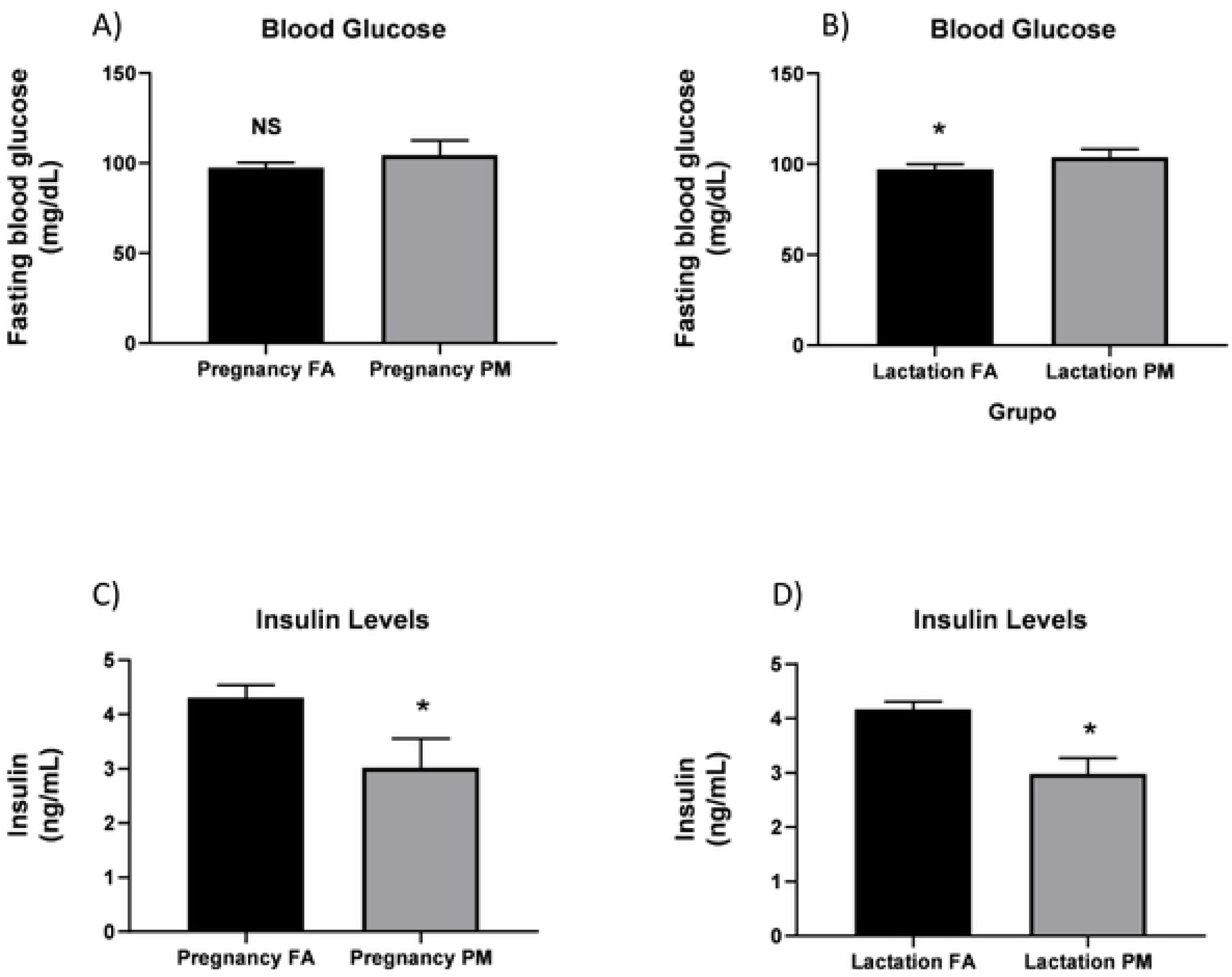
Blood glucose and insulin levels of the offspring (F2) of mice (F1) whose dams (F0) were exposed to PM_2.5_ or FA during either pregnancy or lactation. Values include fasting blood glucose (mg/dL) and serum insulin (ng/mL). Data are shown as mean ± SD (n = 6 mice/group). Mice were 11-week old and fasted overnight before the experiments. A two-tailed Student’s t-test was used for statistical analysis. * *p* < 0.05 vs. FA.

Serum insulin levels were lower in F1 generation mice exposed to air pollution (PM_2.5_) (figure 1, C and D) and remained in generation F2 (figure 2, C and D).

We tested whether the exposure to PM_2.5_ negatively affected glucose homeostasis with an insulin tolerance test. K_ITT_ showed higher insulin resistance among the offspring of mice exposed to FA during lactation (figure 3, A and E), being lower in the offspring of mice exposed to PM_2.5_ during lactation when compared to the control group. Consequently, the AUC values during the ITT of the groups whose mothers were exposed to PM_2.5_ were greater when compared to the control group (figure 3, C and G).

**Figure 3.**
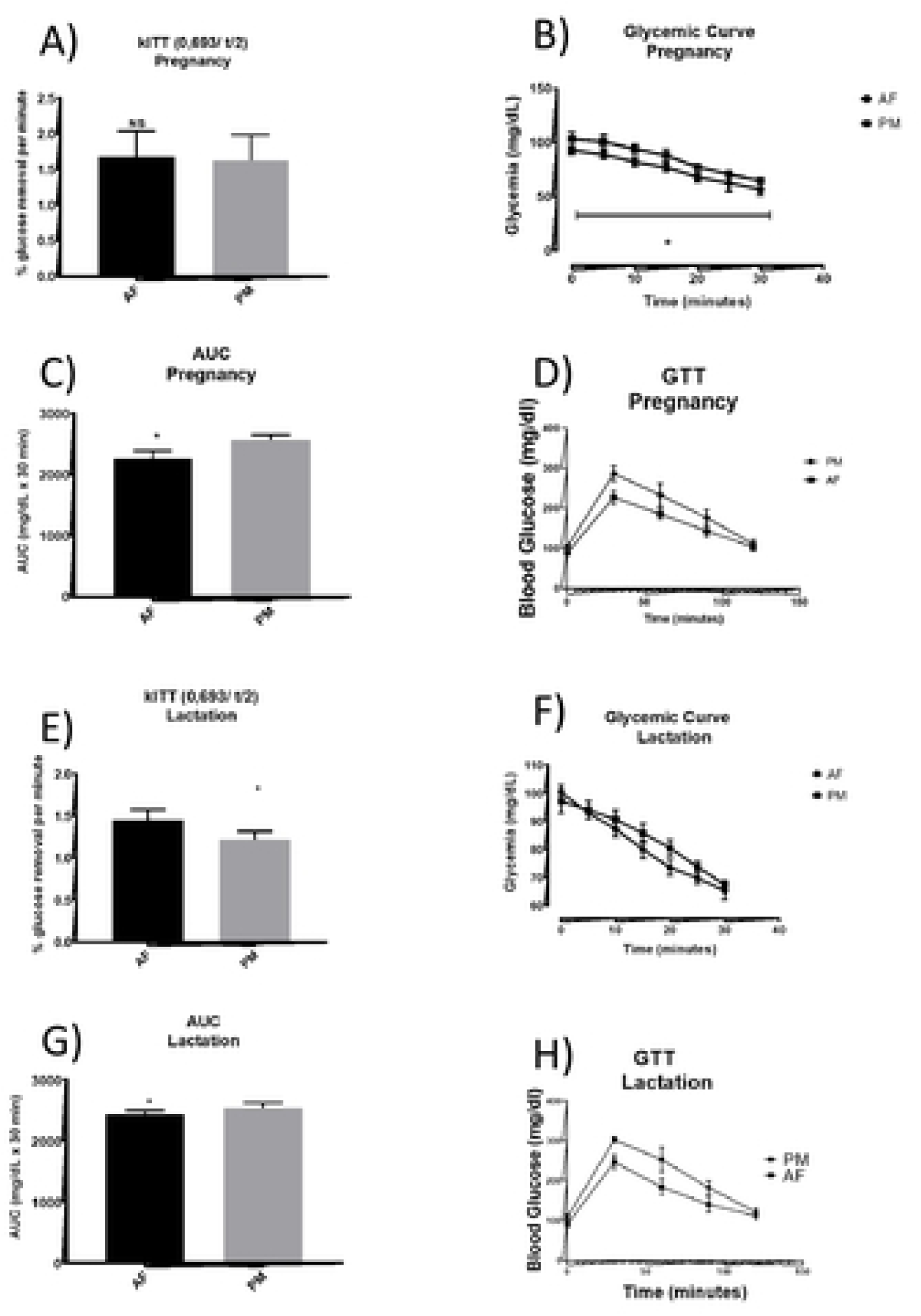
ITT and GTT of the offspring (F1) of mice exposed to PM_2.5_ or FA during either pregnancy or lactation. **A**. Intraperitoneal insulin tolerance test (1.5 insulin unit/kg of body weight) was performed after 8-hour fasting when mice were 11-week old. **B**. Intraperitoneal GTT (2 g of glucose/kg of body weight) was performed in 12-week old mice fasted overnight. Data are shown as mean (n = 6 mice/group) * *p* < 0.05.

During the ITT, blood glucose levels were more elevated among the offspring of mice exposed to PM_2.5_ during pregnancy when compared to the control group throughout the whole test (figure 3B).

As for F2 generation animals, K_ITT_ remained elevated among the control groups (figure 4, A and E). During the ITT, there was no difference in blood glucose levels (figure 4, B and F), therefore no difference in AUC values was observed (figure 4, C and G).

**Figure 4.**
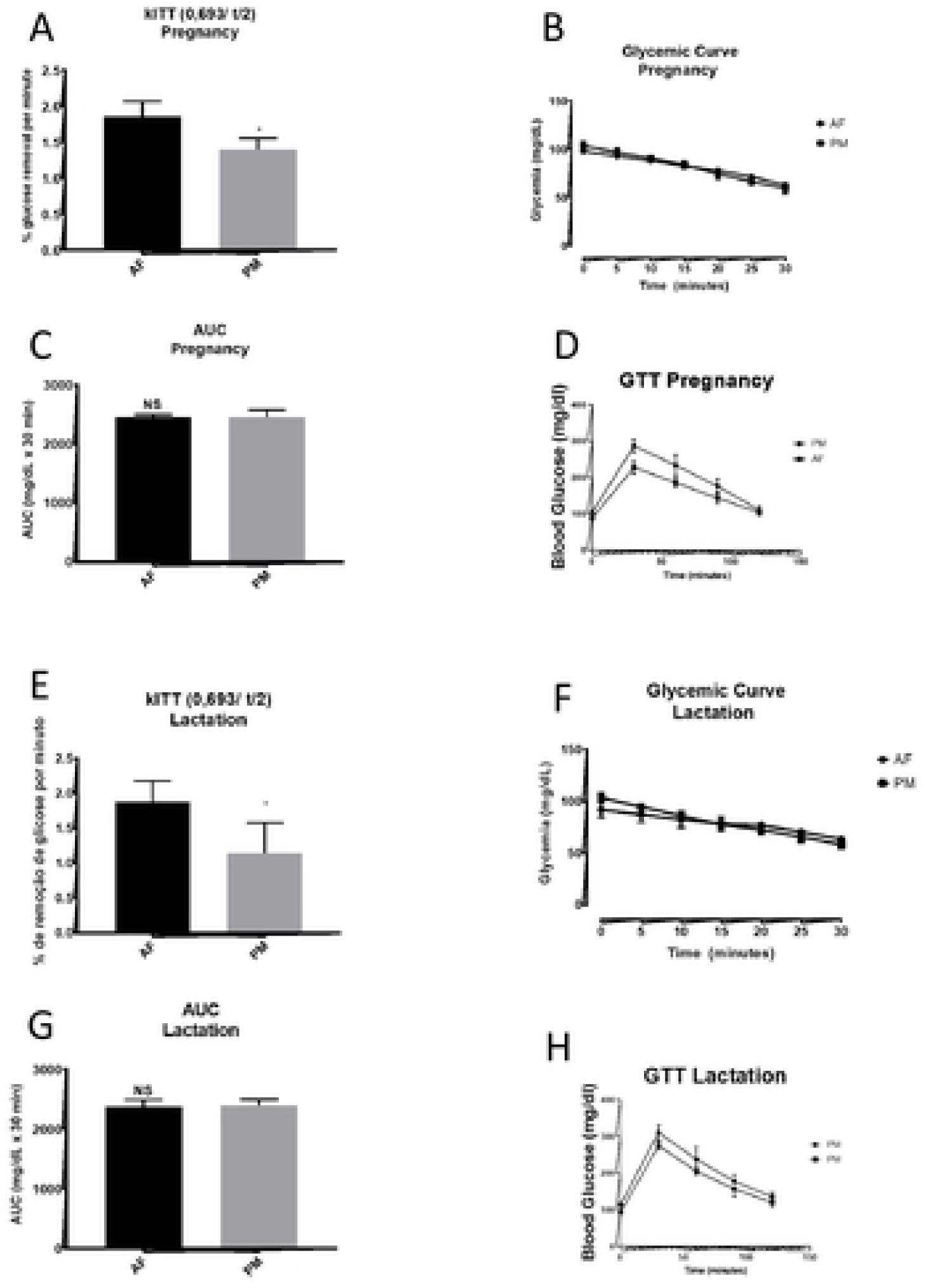
ITT and GTT of the offspring (F2) of mice (F1) whose dams (F0) were exposed to PM_2.5_ or FA during either pregnancy or lactation. **A**. Intraperitoneal insulin tolerance test (1.5 insulin unit/kg of body weight) was performed after 8-hour fasting when mice were 11-week old. **B**. Intraperitoneal GTT (2 g of glucose/kg of body weight) was performed in 12-week old mice fasted overnight. Data are shown as mean (n = 6 mice/group) * *p* < 0.05.

### 2.2. Exposure to PM_2.5_ causes multigenerational alterations in *Pdx1* and *NEUROG3* DNA methylation levels

Considering that the epigenome has been increasingly regarded as an important link between inherited genome changes, we conducted methylation level analyses within the promoter regions of the genes *Pdx1* and *NEUROG3* in the pancreatic islets of F1 and F2 mice.

Our results show that *Pdx1* gene methylation levels were greater among the offspring (F1) of mice exposed to PM_2.5_ during pregnancy (mean = 57.57%) in comparison to the control group (31.46%) (*p* = 0.0001). The lowest methylation levels were identified among the offspring of dams exposed to filtered air during lactation (mean = 18.61%) the mean for the group exposed to PM_2.5_ was 39.01% (*p* = 0.0001) (figure 5A).

**Figure 5.**
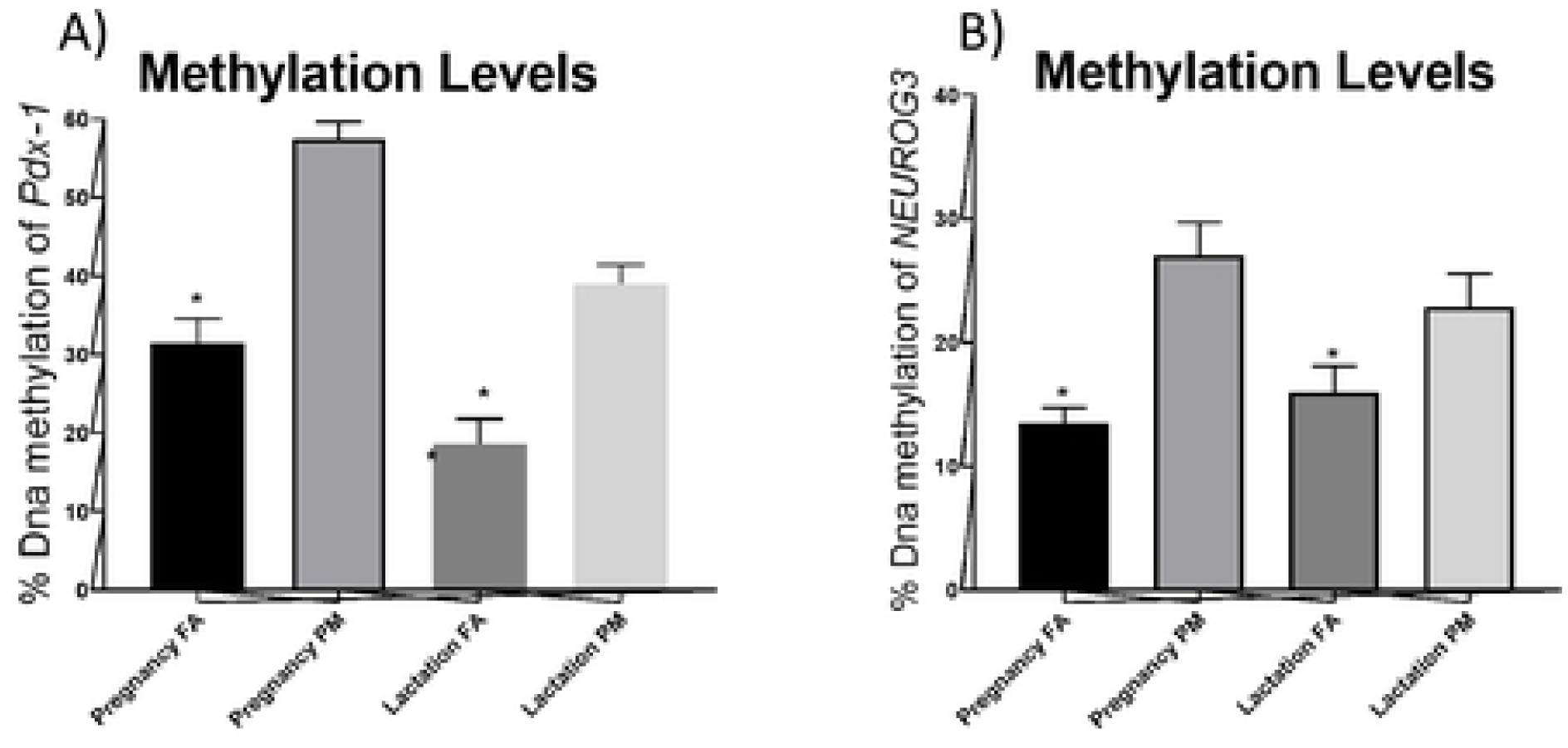
DNA methylation levels of promoter regions within the pancreatic islets of F1 generation animals were analyzed using the HRM method. **A**. *Pdx1* promoter region methylation levels. **B**. *NEUROG3* promoter region methylation levels. Values are expressed as DNA methylation percentages (n = 6 mice/group). * *p* < 0.05.

A similar pattern was observed within the F2 generation, with elevated *Pdx1* methylation levels among the offspring (F2) of mice whose dams were exposed to PM_2.5_ during pregnancy presenting an average of 41.54% and the control group, exposed to filtered air, an average of 23.61% (*p* = 0.004) (figure 6A). In the group exposed during lactation, the PM_2.5_-exposed offspring had a methylation average of 28.24% and the control exposed to filtered air (AF) of 15.94% (*p* = 0.0005).

**Figure 6.**
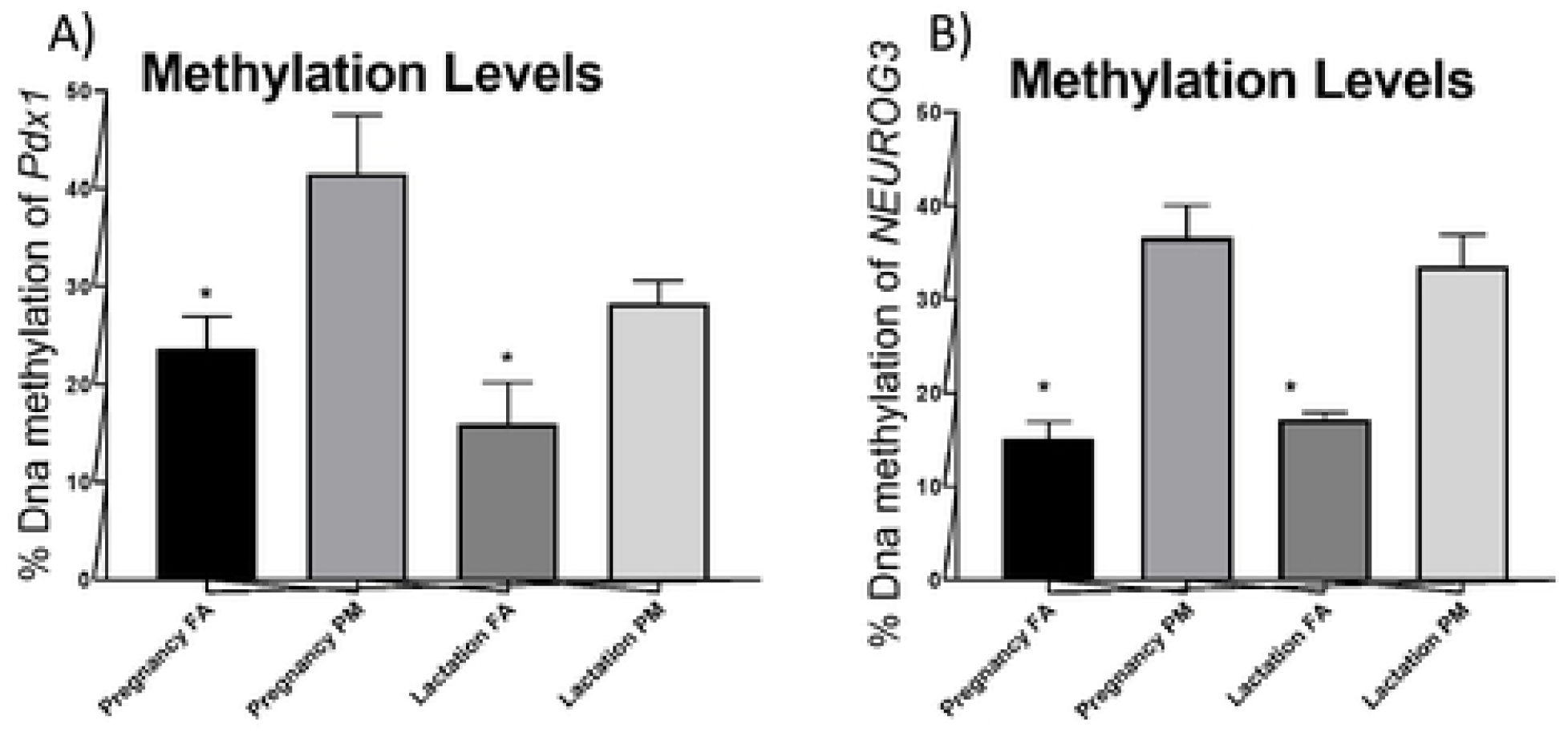
DNA methylation levels of promoter regions within the pancreatic islets of F2 generation animals analyzed using the HRM method. **A**. *Pdx1* promoter region methylation levels. **B**. *NEUROG3* promoter region methylation levels. Values are expressed as DNA methylation percentages (n = 6 mice/group). * *p* < 0.05.

Likewise, to *Pdx1* the *NEUROG3* is a critical transcription factor for β-cell development and maturing. We observed higher DNA methylation levels for that gene within the pancreatic islets of the offspring (F1) of mice exposed to PM_2.5_ during pregnancy (mean = 39.01%) when compared to the control group exposed to filtered air (AF) (mean = 18.61%) (*p* = 0.0001) (figure 5B), in generation (F2) the groups with the highest methylation levels were those exposed to polluted air (PM_2.5_) during pregnancy and lactation (mean = 36.59% and mean = 33.56%), respectively), when compared to the control group (FA) (mean = 15.52%, mean = 17.18%, respectively) (figure 6B).

### 2.3. Impacts of PM_2.5_ exposure on gene expression within mice pancreatic islets

Furthermore, we investigated the impact of exposure to PM_2.5_ on gene expression within pancreatic islets. The offspring (F1) of animals exposed to PM_2.5_ during pregnancy presented lower levels of Pdx1 expression (mean = 2.49,) when compared to the control group (AF) (mean = 7.49) (*p* = 0.0001), in the group exposed during lactation, the mean air polluted group was 7.23 and the control group (AF) 5.17 (*p* = 0.0001) (figure 7A). These lower levels of expression in the pregnant group exposed to PM_2.5_ remained in the next generation (F2) (mean = 3.59) and the control group averaged 6.67 (*p* = 0.0006) (figure 8A).

**Figure 7.**
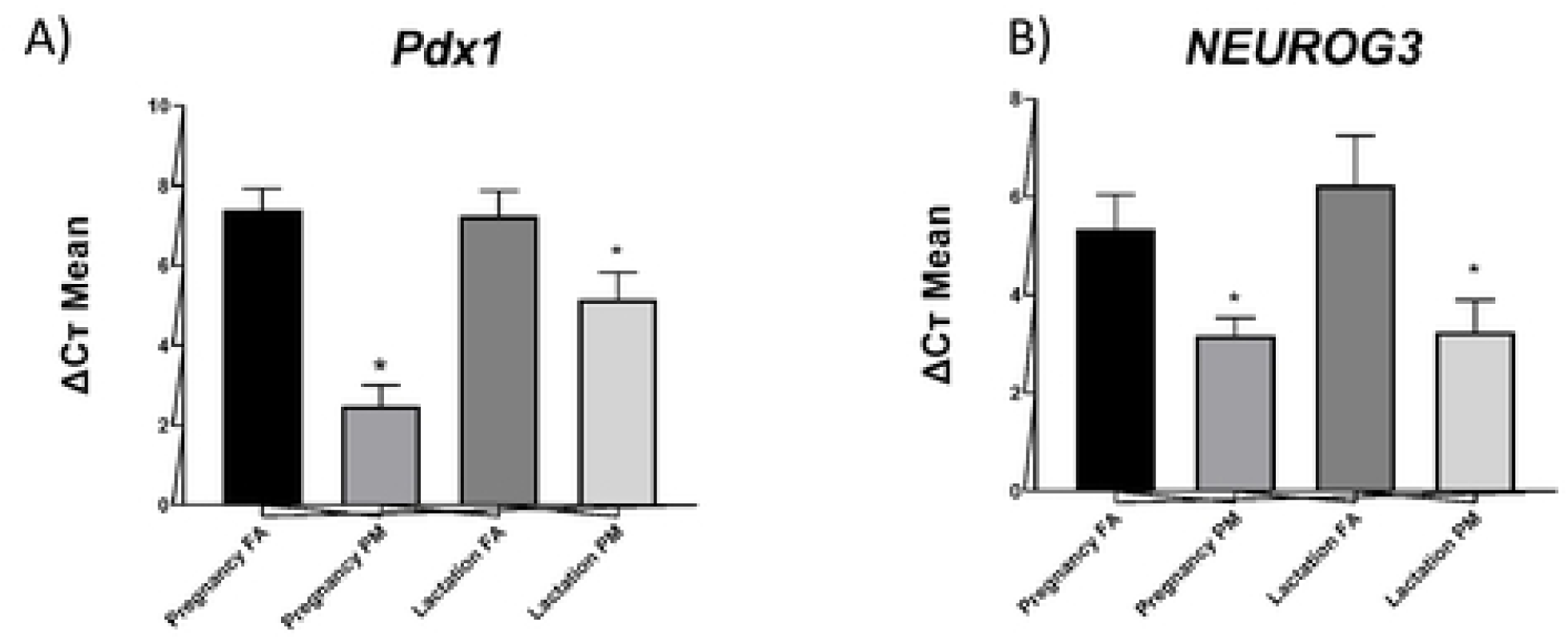
Gene expression levels within pancreatic islets of F1 generation mice. **A**. *Pdx1* gene expression levels. **B**. *NEUROG3* gene expression levels. Data are shown as mean (n = 6 mice/group). * *p* < 0.05.

**Figure 8.**
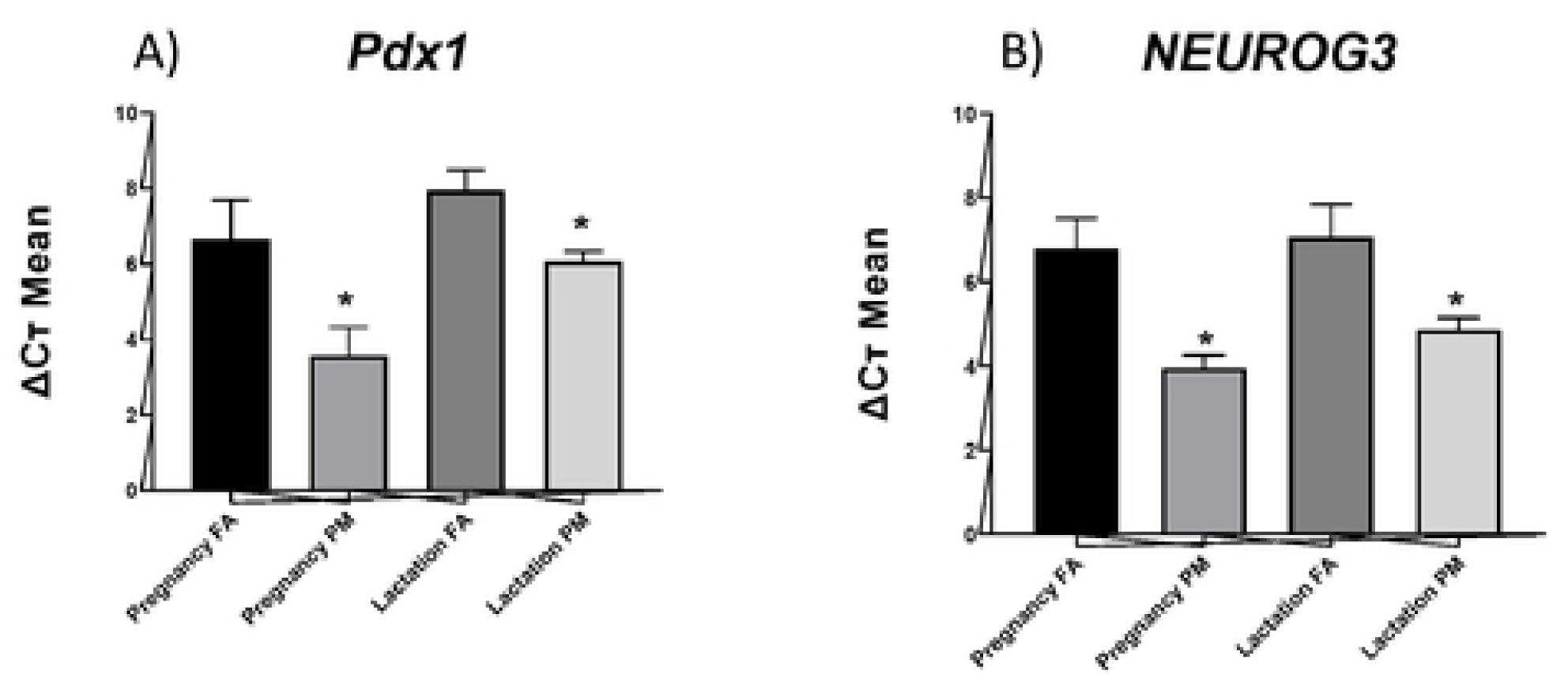
Gene expression levels within pancreatic islets of F2 generation mice. **A**. *Pdx1* gene expression levels. **B**. *NEUROG3* gene expression levels. Data are shown as mean (n = 6 mice/group). * *p* < 0.05.

*NEUROG3* gene expression was also lowered among the offspring (F1) of mice exposed to PM_2.5_ during pregnancy (mean = 3.18) when compared to the control group (AF) (mean =5.34) (*p* = 0.0003) as well as those exposed during lactation to polluted air (PM_2.5_) (mean = 3.25) and filtered air (AF) (mean = 6.25) (*p* = 0.0005) (figure 7B). This decrease was also passed on to the subsequent generation (F2), the mean of exudes exposed to PM_2.5_ during pregnancy was mean = 3.85 and the AF control group with a mean of 6.80 (*p* = 0.0001) in the group exposed during polluted air lactation (PM_2.5_) averaged 4.86 and control (AF) 7.07 (*p* = 0.0003).

## 3. DISCUSSION

This study investigated the impact of F0 generation exposure to PM_2.5_ during either pregnancy or lactation on metabolic parameters as well as on the levels of *Pdx1* and *NEUROG3* gene methylation and gene expression in the two following in F1 and F2 generations.

We demonstrated that exposure to polluted air (PM_2.5_) elevates blood glucose levels in generations F1 and F2. We also showed that DNA methylation levels in the promoter region within the genes *Pdx1* and *NEUROG3* are greater among animals exposed to polluted air during intrauterine development (F1) and that such effect persists for the subsequent generation (F2). F0 exposure to PM_2.5_ during pregnancy was associated with a reduction in gene expression among generations F1 and F2.

These results suggest that the effects caused by exposure to PM_2.5_ during pregnancy and lactation may be permanent, leading to population-wide alterations as a relatively short period of intrauterine exposure to PM_2.5_ may cause persistent gene-specific DNA methylation.

Our results corroborate the findings of Yang et al. [15] which showed increased DNA methylation of the gene *Pdx1* combined with reduced *Pdx1* expression within pancreatic islets of participants diagnosed with type 2 diabetes when compared to a control group. We thus suggest PM_2.5_ exposure as linked with the genesis of type 2 diabetes along with a multigenerational modulation of DNA methylation levels.

DNA methylation is currently the most widely investigated epigenetic mechanism, which consists of the addition of a methyl group to the 5’ position of cytosine residues located in a CG dinucleotide [10]. DNA methylation of gene promoters is generally regarded as a mechanism that represses gene expression [16], which was observed in our study as higher methylation levels of the genes *Pdx1* and *NEUROG3* led to a reduction in the expression of these genes.

Considering that DNA methylation is critical for processes such as embryonic development of mammals, tissue-specific gene expression regulation and genomic imprinting [17], it may suggest that exposure to air pollution during pregnancy and lactation methylated the aforementioned genes, passing such genomic imprinting on to following generations.

*Pdx1* is a pancreatic and duodenal homeobox 1 transcription factor that regulates pancreatic development and β-cell differentiation. It is expressed during the early stage of β-cell development and is crucial to its function [18]. Evidence shows that both genetic and acquired reductions in *Pdx1* expression may lead to type 2 diabetes, β-cell dysfunction, and compromised islets compensation to insulin resistance in human and animal models alike [19– 21].

Adult post-diabetes-onset methylation may cause permanent silencing of *Pdx1* expression. Likewise, an abnormal intrauterine environment may lead to epigenetic alterations that regulate the main genes associated with β-cell development, leaving *Pdx1* expression permanently lowered within β cells [20].

*NEUROG3* is a member of the basic helix-loop-helix transcription factor family involved in neurogenesis and pancreatic embryonic development. In mouse pancreatic embryonic development, *NEUROG3* expression is initially observed in the dorsal pancreatic epithelium at E9.5 and E15.5, later decreasing to substantially lower levels within the neonatal pancreas [22].

Four different types of islet cells (alpha, beta, delta, and pancreatic polypeptide), as well as endocrine precursor cells, were shown as absent in N*EUROG3*-deficient mice, which died postnatally from diabetes [23].

Insulin secretion deficiency underlies the transition from normoglycemia to hyperglycemia in type 1 and type 2 diabetes alike [17]. Though it is known that circulating insulin originates almost exclusively from β cells located in pancreatic islets, our comprehension about the molecular mechanisms responsible for regulating β-cell function in healthy and diseased mammals alike remain incomplete. Pdx1 is a protein that plays a key role in regulating β-cell function [24].

Sun et al. [2] conducted a study with lean C57BL/6 mice exposed to PM_2.5_ for 128 days and didn’t find significant associations between exposure to air pollution and insulin resistant, which may suggest that exposure to air pollution is more deleterious during critical stages of life, such as intrauterine development.

Therefore, our study’s findings show that mice exposed to PM_2.5_ from air pollution during the early life stages are predisposed to developing type 2 diabetes during their lives.

*Pdx1* and *NEUROG3* promoter regions epigenetic modifications are underlying mechanisms that explain the programming effect observed in this study. These alterations may lower insulin levels, thus negatively affecting glucose homeostasis.

Although further research is necessary to elucidate whether the programmed diabetes phenotype can be reprogrammed by interventions posterior to birth and weaning, our results demonstrate that the perinatal development period serves as a critical time window during which mother exposure to PM_2.5_ from polluted air may negatively influence the health of children and grandchildren throughout their lifespan.

Given that air pollution is a major environmental problem, particularly in developing countries and due to poor environmental management and control, these results indicate that harmful epigenetic programming during critical developmental stages is a potential factor associated with the worldwide increase in rates of diabetes.

## 4. METHODS

### 4.1. Animals

C57BL/6J mice were provided by the University of Campinas’ multidisciplinary center of biological research (CEMIB) (University of Campinas, SP, Brazil).

All the experiments conducted in this study complied with the Committee for Ethics in Animal Use of the University of Campinas – CEUA/UNICAMP guidelines, which has approved all of the study’s experimental protocols.

Mice were housed in an animal facility with a room temperature of 22°C, controlled humidity, and a 12-hour light and dark cycle. They were provided with free access to tap water and a standard rodent chow (3.39 Kcal/g; 63.4% calories from carbohydrates, 25.9% calories from protein, and 10.6% calories from fat; Nuvilab CR-1, Nuvital Quimtia, Brazil).

### 4.2. PM_2.5_ or Filtered Air Exposures

Mice were exposed to PM_2.5_ or filtered air (FA) at the same time of the day after one week of acclimation. The Harvard Ambient Fine Particles Concentrator located at the University of São Paulo (USP), in São Paulo, Brazil [25] was used to capture the air and concentrate the PM_2.5_ to obtain a final amount of 600 μg/m3/24 h. Following each exposure, the animals returned immediately to their home cages and were maintained in an animal facility with high-efficiency particulate air (HEPA) filtered air. The duration of the exposure period depended on the ambient concentration of PM_2.5_, which was checked every day in CETESB (Companhia Ambiental do Estado de São Paulo; https://cetesb.sp.gov.br/) and monitored during exposure using a real-time particle sensor; however, the duration never exceeded 3 hours per day.

The specific experimental design was the following: the first set of experiments included six groups of randomly allocated female mice (F0), three of which were exposed to PM_2.5_ throughout pregnancy whereas the other three groups were exposed to FA throughout pregnancy. The second set of experiments included six other groups of randomly allocated female mice, three of which were exposed to PM_2.5_ throughout the lactation period (21 days) whereas the three other groups were exposed to FA throughout the lactation period (21 days).

After the exposition periods, dams were euthanized and offspring (F1) were submitted, 11 weeks after birth, to *in vivo* analyses, after which they were placed for breeding (using three females and one male from each of the 7 litters), being euthanized subsequent to the end of the lactation period. Their body tissues were kept for analyses and their offspring (F2) were submitted to *in vivo* analyses 11 weeks after their birth, after which they were euthanized and had their body tissues analyzed, in accordance with the study’s aims.

### 4.3. Metabolic Parameters

All metabolic parameters were obtained after the exposition. Serum glucose and insulin levels were assessed after an overnight fast.

#### 4.3.1. Insulin Tolerance Test (ITT) and Glucose Tolerance Test (GTT)

In order to perform the ITT, mice were submitted to an 8-hour fasting period and the test began at 8 a.m. Animals received an intraperitoneal (IP) injection of recombinant insulin (Humulin®, Eli Lilly, Indianapolis, IN) at a dose of 1.5 units per kilogram of body weight followed by blood glucose measurements with a glucose meter every 5 minutes up to 30 using blood samples obtained from the animals’ tails.

Glucose disappearance rate (K_ITT_) was obtained using the formula 0.693/t_1/2_ [26]. Blood glucose half-life time (t_1/2_) was estimated from the slope of least square analysis of the blood glucose levels when there was a linear decline [26,27]. The serum glucose area under the curve (AUC) was calculated using the trapezoidal rule [28].

Glucose tolerance test (GTT) was performed after 12 hours of overnight fasting. Mice received IP injections of 2g of glucose per kilogram of body weight. Blood glucose was measured with a glucose meter every 30 minutes up to 120 using blood samples obtained from the animals’ tails.

### 4.4. Tissue collection

Mice were euthanized 12 weeks after birth and had their pancreatic islets collected for further analysis.

#### 4.4.1. Pancreatic islets collection

Mice had their pancreas perfused with Hanks’ balanced salt solution containing 3 mg collagenase/ml and excised immediately after euthanasia. Islets were then dissociated through continuous shaking at 37∼ for 18 min. Isolated islets were then collected [29].

#### 4.4.2. Tissue collection for gene expression by qPCR analysis

Total RNA was extracted from the samples using TRIzol™ Reagent (REF 15596018, Life Technologies, CA, USA) according to the manufacturer’s instructions. A NanoDrop 2000™ spectrophotometer was used to quantify the extracted RNA and 3 µg of total RNA from each sample was used to synthesize a single-strand cDNA utilizing a commercial kit (cat #4368814, High-Capacity cDNA Reverse Transcription Kit, Applied Biosystem, CA, USA), according to the manufacturer’s instructions. A real-time PCR was performed using the QuantStudio 6 Flex Real-Time PCR System (# 4485694, Applied Biosystems, CA, USA) with the following reagents: Universal Master Mix TaqMan™, TaqMan™ Primer (10 µM), cDNA sample (100 ng) and Milli-Q™ H_2_O. The software Data Assist ™ (Applied Biosystems, CA, USA) was used to obtain the cycle limit.

Relative expression levels were determined and then normalized to the expression levels of *GAPDH*. All conditions were performed as duplicates. Primer sequences were ordered from Life Technologies (CA, USA): *Pdx1* - Mm00435565_m1; *NEUROG3* - Mm00437606_s1. Gene transcripts of interest were quantified relative to the internal control *GAPDH* using the 2-^ΔΔCT^ method [30].

#### 4.4.3 Methylation analysis

DNA was isolated using the PureLink™ Genomic DNA Mini Kit (Invitrogen, Thermo Fisher Scientific) according to the protocol provided by the manufacturer.

For the DNA standards and mouse samples, whole DNA was quantified on a NanoDrop 2000™ spectrophotometer (Thermo Scientific, USA). The Cells-to-CpG™ Bisulfite Conversion Kit (Applied Biosystems, Life Technologies, CA, USA) was used according to the manufacturer’s instructions for genomic DNA bisulfite conversion. PCR amplification and High-Resolution Melting (HRM) were conducted using the software Applied Biosystems 7500 Fast System [31].

The final volume of the solution used for PCR was 20 µl, containing: AmpliTaq™ Gold 360 Buffer, 10x, 25mM Mg Chloride, MeltDoctor™ HRM Dye (20x), 25nM of each initiator containing the primer sequences for the genes *Pdx1* (Forward: F-5’-TTTAGGTTAATGATGGTTTTAGGGTAA 3’ and Reverse: 5’ CTCCACTACTCTCCTAAAAAACCAA 3’), *NEUROG3* (Forward: F-5’-TTAGGATGGAGTTAGTTTGTGAAATAT 3’ and Reverse: 5’

CATAAAATCTTTTAACTCAAAAAAAA3’), and 1 µl of bisulfite-converted genomic DNA. The qPCR consisted of an initial enzyme activation at 95 °C which lasted for 10 minutes, followed by 40 cycles, each consisting of 15 s at 95 °C, 60 s at 60 °C, 10 s at 95 °C, 60 s at 60 °C, 15 s at 95 °C, and 15 s at 60 °C. For *Pdx1*, the 148-bp amplification included a total of 4 CpG dinucleotides between the primers and ranged from 91 to 177 in the promoter region. For *NEUROG3*, the 170-bp amplification included a total of 9 CpG dinucleotides between the primers and ranged from 85 to 111 in the promoter region.

As positive (100% methylated) and negative (0% methylated) controls, we used CpGenome™ Universal Methylated and Unmethylated DNA (Chemicon, Millipore, Billerica, MA, USA), respectively. All reactions were performed at least in triplicate.

### 4.5. Assays

Serum insulin concentration was measured using a commercial ELISA kit (Abcam’s #EZRMI-13K, Cambridge, UK). The enzymatic colorimetric method was used to measure serum glucose.

### 4.6. Statistical analysis

GraphPad Prism software (San Diego, CA, USA) was used to calculate the mean and standard deviation (SD), as well as to perform statistical analyses and to produce the graphics. All the results were expressed as means ± SD. One-way ANOVA or two-way ANOVA with post hoc test (Tukey), and an unpaired two-tailed Student’s t-test were applied in which p < 0.05 determined the significant difference.

## REFERENCES

1. Rajper SA, Ullah S, Li Z. Exposure to air pollution and self-reported effects on Chinese students: A case study of 13 megacities. PLoS One. 2018;13: 1–21. doi:10.1371/journal.pone.0194364

2. Sun Q, Yue P, Deiuliis JA, Lumeng CN, Kampfrath T, Mikolaj MB, et al. Ambient Air Pollution Exaggerates Adipose Inflammation and Insulin Resistance in a Mouse Model of Diet-Induced Obesity. Circulation. 2009;119: 538–546. doi:10.1161/CIRCULATIONAHA.108.799015

3. Chen M, Qin X, Qiu L, Chen S, Zhou H, Xu Y, et al. Concentrated ambient PM2.5 - induced inflammation and endothelial dysfunction in a murine model of neural IKK2 deficiency. Environ Health Perspect. 2018;126: 1–10. doi:10.1289/EHP2311

4. Liang F, Yang X, Liu F, Li J, Xiao Q, Chen J, et al. Long-term exposure to ambient fine particulate matter and incidence of diabetes in China: A cohort study. Environ Int. 2019;126: 568–575. doi:10.1016/j.envint.2019.02.069

5. Sun Z, Zhu D. Exposure to outdoor air pollution and its human health outcomes: A scoping review. PLoS One. 2019;14: 1–18. doi:10.1371/journal.pone.0216550

6. Shen GF, Yuan SY, Xie YN, Xia SJ, Li L, Yao YK, et al. Ambient levels and temporal variations of PM2.5 and PM 10 at a residential site in the mega-city, Nanjing, in the western Yangtze River Delta, China. J Environ Sci Heal - Part A Toxic/Hazardous Subst Environ Eng. 2014;49: 171–178. doi:10.1080/10934529.2013.838851

7. Alfano R, Herceg Z, Nawrot TS, Chadeau-Hyam M, Ghantous A, Plusquin M. The Impact of Air Pollution on Our Epigenome: How Far Is the Evidence? (A Systematic Review). Curr Environ Heal Reports. 2018;5: 544–578. doi:10.1007/s40572-018-0218-8

8. Saino N, Albetti B, Ambrosini R, Caprioli M, Costanzo A, Mariani J, et al. Inter-generational resemblance of methylation levels at circadian genes and associations with phenology in the barn swallow. Sci Rep. 2019;9: 6505. doi:10.1038/s41598-019-42798-3

9. de F.C. Lichtenfels AJ, van der Plaat DA, de Jong K, van Diemen CC, Postma DS, Nedeljkovic I, et al. Long-term Air Pollution Exposure, Genome-wide DNA Methylation and Lung Function in the LifeLines Cohort Study. Environ Health Perspect. 2018;126: 027004. doi:10.1289/EHP2045

10. Obri A, Claret M. The role of epigenetics in hypothalamic energy balance control: implications for obesity. Cell Stress. 2019;3: 208–220. doi:10.15698/cst2019.07.191

11. Lee MK, Xu CJ, Carnes MU, Nichols CE, Ward JM, Kwon SO, et al. Genome-wide DNA methylation and long-term ambient air pollution exposure in Korean adults. Clin Epigenetics. 2019;11: 1–12. doi:10.1186/s13148-019-0635-z

12. Glavas MM, Hui Q, Tudurí E, Erener S, Kasteel NL, Johnson JD, et al. Early overnutrition reduces Pdx1 expression and induces β cell failure in Swiss Webster mice. Sci Rep. 2019;9: 3619. doi:10.1038/s41598-019-39177-3

13. Daxinger L, Whitelaw E. Understanding transgenerational epigenetic inheritance via the gametes in mammals. Nat Rev Genet. 2012;13: 153–162. doi:10.1038/nrg3188

14. Skinner MK. What is an epigenetic transgenerational phenotype? Reprod Toxicol. 2008;25: 2–6. doi:10.1016/j.reprotox.2007.09.001

15. Yang BT, Dayeh TA, Volkov PA, Kirkpatrick CL, Malmgren S, Jing X. Aumento da metilação do DNA e diminuição da expressão de PDX-1 em ilhotas pancreáticas de pacientes com diabetes tipo 2 Abstrato. 2019;26: 1203–1212.

16. Moore LD, L. T, Fan G. DNA Methylation and Its Basic Function. Neuropsychopharmacology. 2013;38: 23–38. doi:10.1038/npp.2012.112

17. Bansal A, Pinney SE. DNA methylation and its role in the pathogenesis of diabetes. Pediatr Diabetes. 2017;18: 167–177. doi:10.1111/pedi.12521

18. Ryu GR, Lee E, Kim JJ, Moon SD, Ko SH, Ahn YB, et al. Comparison of enteroendocrine cells and pancreatic β-cells using gene expression profiling and insulin gene methylation. Nishimura W, editor. PLoS One. 2018;13: e0206401. doi:10.1371/journal.pone.0206401

19. Park JH, Stoffers DA, Nicholls RD, Simmons RA. Development of type 2 diabetes following intrauterine growth retardation in rats is associated with progressive epigenetic silencing of Pdx1. J Clin Invest. 2008;118: 2316–2324. doi:10.1172/JCI33655

20. Wang L, Fan H, Zhou L, Wu Y, Lu H, Luo J. Altered expression of PGC-1 α and PDX1 and their methylation status are associated with fetal glucose metabolism in gestational diabetes mellitus. Biochem Biophys Res Commun. 2018;501: 300–306. doi:10.1016/j.bbrc.2018.05.010

21. Gao T, McKenna B, Li C, Reichert M, Nguyen J, Singh T, et al. Pdx1 Maintains β Cell Identity and Function by Repressing an α Cell Program. Cell Metab. 2014;19: 259–271. doi:10.1016/j.cmet.2013.12.002

22. Apelqvist Å, Li H, Sommer L, Beatus P, Anderson DJ, Honjo T, et al. Notch signalling controls pancreatic cell differentiation. Nature. 1999;400: 877–881. doi:10.1038/23716

23. Zhu Y, Liu Q, Zhou Z, Ikeda Y. PDX1, Neurogenin-3, and MAFA: Critical transcription regulators for beta cell development and regeneration. Stem Cell Research and Therapy. Stem Cell Research & Therapy; 2017. pp. 1–7. doi:10.1186/s13287-017-0694-z

24. Maganti A V, Maier B, Tersey SA, Sampley ML, Mosley AL, Özcan S, et al. Transcriptional activity of the islet β cell factor Pdx1 Is augmented by lysine methylation catalyzed by the methyltransferase Set7/9. J Biol Chem. 2015;290: 9812–9822. doi:10.1074/jbc.M114.616219

25. Ribeiro AACM, Dolhnikoff M, Caldini EG, Damaceno-Rodrigues NR, Veras MM, Saldiva PHN, et al. Particulate Urban Air Pollution Affects the Functional Morphology of Mouse Placenta1. Biol Reprod. 2008;79: 578–584. doi:10.1095/biolreprod.108.069591

26. Bonora E, Moghetti P, Zancanaro C, Cigolini M, Querena M, Cacciatori V, et al. Estimates of In Vivo Insulin Action in Man: Comparison of Insulin Tolerance Tests with Euglycemic and Hyperglycemic Glucose Clamp Studies*. J Clin Endocrinol Metab. 1989;68: 374–378. doi:10.1210/jcem-68-2-374

27. Kuga GK, Muñoz VR, Gaspar RC, Nakandakari SCBR, da Silva ASR, Botezelli JD, et al. Impaired insulin signaling and spatial learning in middle-aged rats: The role of PTP1B. Exp Gerontol. 2018;104: 66–71. doi:10.1016/j.exger.2018.02.005

28. Matthews JNS, Altman D, Campbell MJ, Royston P. Analysis of serial measurements in medical research: Authors’ reply. BMJ. 1990;300: 680–680. doi:10.1136/bmj.300.6725.680-a

29. Teixeira CJ, Santos-Silva JC, de Souza DN, Rafacho A, Anhe GF, Bordin S. Dexamethasone during pregnancy impairs maternal pancreatic β-cell renewal during lactation. Endocr Connect. 2019;8: 120–131. doi:10.1530/EC-18-0505

30. Bustin SA, Benes V, Garson JA, Hellemans J, Huggett J, Kubista M, et al. The MIQE Guidelines: Minimum Information for Publication of Quantitative Real-Time PCR Experiments. Clin Chem. 2009;55: 611–622. doi:10.1373/clinchem.2008.112797

31. Lima RPA, do Nascimento RAF, Luna RCP, Persuhn DC, da Silva AS, da Conceição Rodrigues Gonçalves M, et al. Effect of a diet containing folate and hazelnut oil capsule on the methylation level of the ADRB3 gene, lipid profile and oxidative stress in overweight or obese women. Clin Epigenetics. 2017;9: 110. doi:10.1186/s13148-017-0407-6

